# Single cell analysis of ovarian immune cells during homeostasis and hormonal flux reveals dynamic changes in NK and B cell populations in the periovulatory phase

**DOI:** 10.1101/2025.04.18.649580

**Authors:** Tia Y. Brodeur, Toby B. Lanser, Lucy Salter, Katherine Sessions, Tyler Hogan, Elizabeth A. Bonney, Elizabeth McGee, Seth Frietze, Dimitry N. Krementsov

**Affiliations:** Indiana University School of Medicine, Department of Obstetrics and Gynecology, Division of Reproductive Endocrinology, Indianapolis, Indiana; Ann Romney Center for Neurologic Diseases, Harvard Medical School, Brigham and Women’s Hospital, Boston, MA; Department of Obstetrics and Gynecology and Reproductive Sciences, University of Vermont, Burlington, Vermont; Department of Biomedical and Health Sciences, University of Vermont, Burlington, Vermont

## Abstract

The mammalian ovary is the dynamic end-organ of the hypothalamic pituitary ovarian axis. In this coordinated system, ovarian cells undergo continuous cycles of apoptosis, proliferation, and differentiation. These changes parallel fluctuations in ovarian hormones such as estradiol (E2); however, the ovarian immune microenvironment during high-E2 and low-E2 states is not fully understood. We induced a high-E2 state in the mouse ovary by stimulation via gonadotropin treatment. Single-cell RNA sequencing and flow cytometry analysis of ovarian leukocytes revealed abundant mature NK cells, B1 and B2 cells, CD8+ T cells, CD4+ T cells, mature CD4-CD8-T cells, T regulatory cells, and distinct myeloid subsets, including *Trem2+* and *Apoe+* macrophages. *In vivo* labeling of circulating cells determined that the vast majority of ovarian leukocytes were tissue-resident. Following gonadotropin treatment, the frequency of NK cells increased two-fold, while B1 cell frequency was reduced by half. Consistently, flow cytometric analysis revealed an increase in mature CD11b+ NK cells following gonadotropin treatment. Gonadotropin treatment also increased cell-cell signaling by myeloid cells at the expense of NK cells. Our findings reveal a diverse resident immune landscape in the ovary that responds robustly to hormonal changes. These findings have implications for a role for immune cells in ovarian physiology and functional dysregulation.

## Introduction

The ovary contains numerous leukocytes, and the processes of development of ovarian follicles and ovulation involves a local immune response. However, if these complex and dynamic processes are not properly regulated, they can result in chronic inflammation and autoimmunity(1). For example, macrophage-deficient *Csf1r-/-* mice and *Csf1^op/op^-*mutant osteopetrotic mice have decreased fertility and this may be in part due to impaired tissue remodeling mediated by macrophages that allows for downstream follicular growth (2) Despite the critical role of ovarian follicular macrophages and other ovary-resident leukocytes; their functional phenotypes and possible regulation by estrogen throughout the estrous cycle remain poorly understood. A mechanistic understanding of immune regulation within the ovary could provide insight into the pathogenesis of diseases such as polycystic ovary syndrome, ovarian endometriosis, and autoimmune primary ovarian insufficiency (POI). Despite its significant clinical impact, including infertility (3), osteoporosis (4, 5), and depression (6), POI remains poorly understood. POI can arise from many causes, including genetic, environmental, metabolic, viral, and autoimmune mechanisms (3, 7, 8). Among these, POI resulting from autoimmunity presents a unique therapeutic challenge, as there is no disease modifying-treatment, despite progress in managing other autoimmune conditions. Estrogen may contribute to autoimmune POI, having been implicated as a driver of autoimmunity (9–11) and potentially explaining the increased female predominance seen in diseases like systemic lupus erythematosus, with one mechanism being lowering the threshold for autoreactivity in B cells (9, 11–14). Moreover, unlike natural menopause or other forms of POI, patients with autoimmune POI can be menopausal due to lymphocytic oophoritis despite oocytes remaining in their ovaries (15). Research on the immune cell phenotypes within the ovary, and their regulation by estrogens has been limited, leaving key mechanisms underlying disease pathogenesis largely unexplored.

Autoimmune POI patients are prone to development of autoantibodies targeting the thyroid and adrenal glands, which can be associated with hypothyroidism and adrenal insufficiency (15–17). These patients have also been shown to have ovarian autoantibodies and oophoritis, although the antibodies have not been established as pathogenic and may instead represent epiphenomena (18). The initiating events leading to autoantibody production are unknown, and the role of ovarian resident B cells has not been established.

Natural killer (NK) cells are cytotoxic innate lymphocytes involved in cancer surveillance, anti-tumor cytotoxicity and antiviral immunity (19, 20). These crucial effector functions are mediated by proinflammatory cytokines, chemokines and activating surface receptors that trigger cytotoxicity via secreted cell-perforating proteins and death receptor engagement on target cells (19, 20). Although ovary-trophic viruses are relatively rare, NK cells have been shown to protect ovarian follicles following MCMV infection through cytokine-mediated immune responses, thereby preserving fertility (21). NK cell-based immunotherapies have been explored in ovarian cancer, yet the role of NK cells in maintaining ovarian homeostasis remains unknown.

## Materials and Methods

### Mice and superovulation

The animal experiments described in this paper were performed under a protocol approved by the University of Vermont Animal Care and Use Committee. Female C57BL/6J mice were purchased at 6-8 weeks from the Jackson Laboratory (Bar Harbor, ME, USA) and treated at 10-12 weeks of age. For superovulation experiments, mice were treated by intraperitoneal (i.p.) injection with 10U of human menopausal gonadotropins (GN, Ferring) or saline for controls. Mice were sacrificed 48 hours after treatment. In some *e*xperiments, ovulation was triggered with human chorionic gonadotropin 5U (hCG) by i.p. injection (Ferring) and mice were sacrificed 24 hours later. In experiments where ovulation was triggered, GN dosing was staggered such that each experimental group would be terminated at the same time. For intravascular labelling, mice were injected with 3μg anti-CD45-PE (BioLegend) as previously described (22) administered intravenously via tail vein or retro-orbitally, and sacrificed after 3 minutes. The mice were maintained on a 12-hour light/dark cycle and had access to food and water ad libitum. Animals were euthanized using CO_2_ followed by cervical dislocation.

### Ovarian leukocyte isolation and flow cytometry

Ovaries were dissected from mice and placed in cold RPMI with 2% fetal bovine serum (FBS, GIBCO) in Petri dishes. 25 gauge needles we used to disperse ovarian cells. Any remaining fibrous tissue was gently and briefly mechanically disrupted with frosted glass slides. Cells were then subjected to Fixable Live/Dead staining (Invitrogen) per manufacturer’s protocol. Cells were treated with diluted Fc receptor blocking antibody (BioLegend) for 20 minutes on ice, pelleted, then stained with fluorescently-conjugated primary antibodies diluted in PBS + 2% FBS as indicated in figures. Samples were washed and resuspended in PBS + 2% FBS and then analyzed by flow cytometry on a Cytek Aurora spectral flow cytometer.

### Single-cell RNA sequencing

Mice were treated i.p. with 10U of human menopausal gonadotropins (Ferring, n=5) or saline for controls (n=5). Mice were sacrificed 48 hours after treatment. Ovary cell suspensions were prepared as described above. Individual mouse samples were stained with unique TotalSeq-B Hashtag antibodies targeting MHC class I and CD45 (BioLegend) to allow for sample multiplexing to maximize cell recovery and minimize batch effects. Cells were subsequently stained with Fixable Live/Dead staining (Invitrogen) per manufacturer’s protocol, followed by fluorescently-labelled antibodies against CD45, CD4, CD19, CD4, and CD8. CD45 positive viable cells were sorted using a FACSAria. Sorted cells were processed by single cell RNA-sequencing (scRNA-seq) using the 10x Genomics Chromium platform and standard protocols, as described in our previous studies (23). Gonadotropin-treated and control samples were loaded on to separate 10x Genomics Chromium flow cells, followed by GEM generation and cell barcoding. Single-cell cDNA libraries were constructed using the Chromium Single Cell 3’ v3 Kit (10x Genomics, USA), with 16 cycles of PCR amplification, per manufacturer’s recommendation. Libraries were pooled equimolarly and sequenced on an Illumina HiSeq 2500 platform (USA). Read alignment, initial quality control, and sample demultiplexing were performed using the *CellRanger* pipeline (10x Genomics), followed by downstream analysis in *Seurat* v5.1 implemented in R, essentially as in our previous studies (23), combining both treatment groups for the initial analysis, and segregating by treatment group for subsequent analyses. For clustering analysis, data dimensionality was assessed using the *JackStrawPlot* and *ElbowPlot* functions. Initial clustering was performed using 20 dimensions. Subsequent sub-clustering was done using 15 dimensions. A small cluster identified as red blood cells was manually removed from the final dataset. Differential gene expression within clusters was assessed using *Seurat’s* Wilcoxon rank-sum test. P-values were adjusted for multiple testing using the Bonferroni or Benjamini-Hochberg correction.

#### CellChat analysis of scRNA-seq data

*Seurat* objects containing annotated cell clusters were converted into *CellChat* objects, and processed using the standard *CellChat v2* pipeline, using the Secreted Signaling mouse database (24). Communication probabilities were inferred using the triMean method. Specific visualization functions used are noted in the corresponding figure legends. For comparative analysis between gonadotropin-treated and control groups, separate *CellChat* objects were generated for each condition. These were subsequently merged for differential signaling analysis.

### Integration of splenocyte scRNA-seq datasets

Integration of ovary and spleen data was performed using *scANVI* v1.2.1, implemented in Python (25). The top 2000 variable genes were selected for integration. The variational autoencoder model was trained using 20 epochs with 2 hidden layers and 30 latent features. Pathway enrichment analysis of tissue-specific differentially abundant genes was performed *gseapy* v1.1.1 (26).

### Immunofluorescence

Ovaries were dissected from mice and frozen in Tissue-Tek O.C.T. (Sakura) atop a dry ice/liquid nitrogen slurry. Ovaries were cryosectioned at 8-micron thickness and adhered to glass slides. Slides were dried overnight, fixed in ice-cold acetone for 10 minutes at −20°C, washed in phosphate buffered saline (PBS) and blocked with PBS + 1% bovine serum albumin (BSA, Sigma) for 1 hour. Primary antibody was diluted in blocking solution and incubated overnight at 4°C. Tissue sections were stained with cleaved caspase-3 (Cell Signaling Technology) or F4/80 (eBioscience). The slides were rinsed three times in PBS, then incubated with fluorescently-labeled secondary antibody in blocking solution for 1-2 hours at room temperature. The slides were then washed three times in PBS and coverslipped with VectaShield Vibrance Antifade Mounting Medium with DAPI (Vector Laboratories). Images were obtained on an Olympus BX50 or Keyence.

### Statistical Methods

For flow cytometry and *in vitro* experiments, statistical analysis was performed using Prism software (version 9, GraphPad). Two-tailed Student’s t-tests were used, with additional details provided in the figure legends.

## Results

### The ovarian immune cell milieu is diverse and includes B cells, T cells, NK cells, and myeloid cells

The ovary is a tissue that undergoes rapid and dynamic structural changes throughout the estrous cycle. Given the role of the immune system in tissue remodeling, we sought to characterize the ovarian immune cell landscape during homeostasis and preovulatory state. To this end, we performed single-cell transcriptomic profiling of ovarian immune cells. Female C57BL/6 (B6) mice were injected with gonadotropin (GN) or PBS control (n=5 per treatment group), followed by ovarian tissue dissociation and isolation of viable CD45^+^ cells by FACS (**Fig. 0, see Methods**). For our initial analysis, to identify and annotate the major ovarian immune cell populations, we combined the data from control and GN-treated mice.

**Figure 0.**
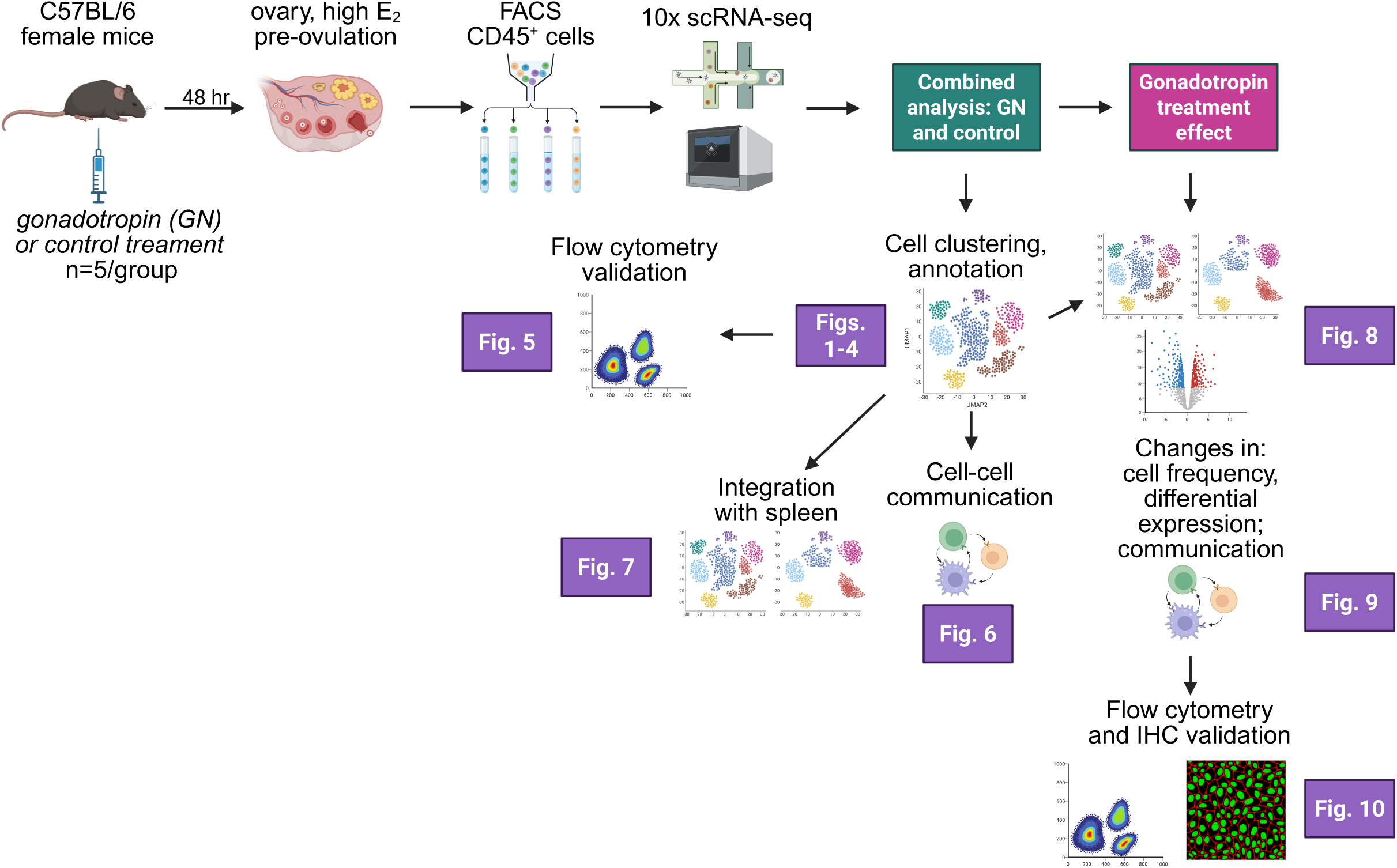
Graphical abstract and study design.

Unsupervised clustering identified 14 major immune cell populations (**Fig. 1A**). Cluster annotation was performed manually, using canonical marker genes (**Fig. 1B**; **Table S1**). The most abundant clusters were three distinct *Cd19+* B cell clusters (B0, B1, and B2). The B2 subset expressed markers of naive B2 cells, including *Fcer2* (encoding CD23), *Ighd* and *Ighm* (encoding IgM and IgD), while the B1 cluster expressed markers associated with the innate-like B1 cells, including *Bhlhe41*, *Zbtb32*, and *Ighm,* in the absence *Ighd* (*27*) (**Fig. 1C**). The intermediate B0 cluster resembled B2 cells but with lower *Ighd* and *Fcer2a* expression, suggesting a less differentiated state. None of the B cell clusters expressed class-switched immunoglobulins (*Ighg1*, *Ighg2a*, *Ighg2c*, *Igha*, and *Ighe* (data not shown)), indicating they were likely naive B cells. The next largest cluster consisted of NK cells, marked by high expression *Ncr1*, *Gzma*, *Nkg7*, and Ly49 family genes (**Fig. 1D, Table S1**). The next most numerous were three T cell clusters, consisted of aβ CD4 T cells (*Cd3e*+, *Trac+*, *Cd4*+), CD8 T cells (*Cd3e*+, *Trac+*, *Cd8b1*+), and a heterogenous population we designated as double-negative (DN) aβ T cells. This DN population showed low expression of *Cd4*, *Cd8a*/*Cd8b1*, and γδ TCR genes (e.g. *Tcrg-C4*), but high expression of *Cd3e* and *Trac* (encoding T cell receptor alpha constant segment), consistent with an aβ TCR+ DN phenotype. Several clusters of myeloid cells were identified (marked by *Lyz2* expression), including neutrophils/PMN (*S100a8*+, *S100a9*+, *Lcn2*+) and tissue-resident macrophages (*Csf1r*, *Adgre4*+) (**Fig. 1B** and **D**). Two additional macrophage clusters were identified, one marked by high expression of *Mertk, Lpar1,* and *Fgfr1* (Mertk+ macs), and another marked by high expression of *C1q* genes (*C1qa, C1qb, C1qc*), *Trem2,* and *Apoe* (C1q+ macs) (**Fig. 1D**). Interestingly, both tissue macs and Mertk+ macs expressed the *Ccr2* receptor typical of monocytes, but Mertk+ macs lacked the monocyte marker *Ly6c2,* while tissue macs clearly expressed tissue resident markers like *Csf1r*, *Adgre4*, *Cx3cr1*, and others (**Fig. 1D** and **Table S1**), suggesting that none of the myeloid populations in the ovary are typical monocytes. Interestingly, C1q+ and Mertk+ macrophages expressed intermediate and high levels, respectively, of MHC class II molecules (*H2-Ab1*, *H2-Aa*; **Fig. 1D** and **Table S1**), suggesting a role in antigen presentation. Minor cell clusters (<100 cells total per cluster) included γδTCR+ NK-like cells (gd-NK; *Tcrg-C4+, Tbx21+, Trac-low*), which surprisingly also expressed the NK receptor *Ncr1,* γδ type 17 cells (gdT17; *Tcrg-C4+, Il23r+, Trac-low*), as well as dendritic cells (DCs; marked by *Siglech* expression and thus most likely corresponding to plasmacytoid DCs) (**Fig. 1B** and **Table S1**). Taken together, these findings reveal a diverse population of immune cells in the ovary, including substantial numbers of B cells, T cells, and NK cells, as well as several phenotypically distinct myeloid subsets.

**Figure 1.**
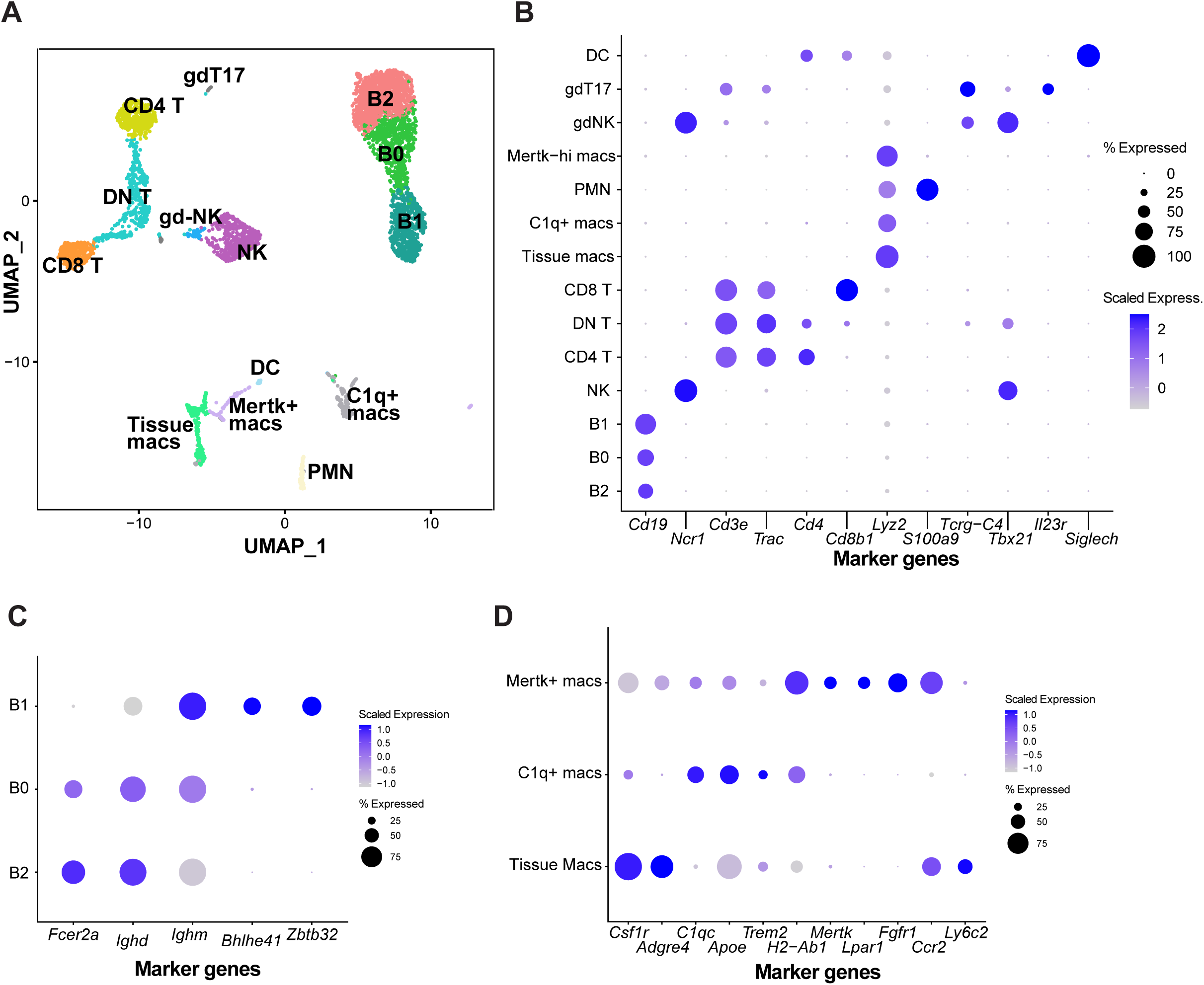
Single cell transcriptional analysis of the ovarian immune landscape. Female C57BL/6 mice were injected with PBS or human menopausal gonadotropin (GN). 48 hours later, ovarian cells were labeled for cell surface markers and CD45+ live cells were isolated using FACS. Isolated cells were pooled by treatment and processed for scRNAseq, and analyzed using the Seurat package, as detailed in the Materials and Methods. Data here represent pooled data for all cells isolated from control and gonadotropin treated mice. (**A**) UMAP-based two-dimensional projection of cell clustering of major cell populations and transcriptional states. (**B**) Dot plot representing scaled expression of selected marker genes for major cell populations. (**C**) Dot plot of scaled expression of selected marker genes for B cell populations. (**D**) Dot plot of expression of selected marker genes for myeloid cell populations.

### Analysis of ovarian B cell subsets identifies B2 and B1a and B1b cells

To further characterize ovarian immune cell subsets, we performed sub-clustering analysis of the three most numerous cell populations: B cells, NK cells, and αβ T cells. For B cells, B0, B1, and B2 populations were selected for re-clustering, and the resulting new clusters were annotated using marker gene identification and manual curation. This analysis revealed a total of seven B cell clusters (**Fig. 2A**). The two largest clusters consisted of B2 and B0 cells, marked by expression of *Fcer2a*, *Ighd*, and *Ighm* (**Fig. 2A-C**). The next two largest clusters we comprised of B1 cells, as marked by *Zbtb32*, *Bhlhe41*, *Ighm*, and lack of *Ighd* and *Fcer2a* (**Fig. 2A-C)**. These were further classified into B1a and B1b cells, as marked by the expression or absence of *Cd5* and *Ctla4*, respectively (**Fig. 2C**, and **Table S2**). The B1b cluster uniquely expressed a high level of *Lifr*, encoding the receptor for LIF) (**Fig. 2C**). Additional minor B cell clusters were identified. These included immature B2 cells (B-imm), marked by expression of *Vpreb3* (encoding the B cell receptor surrogate light chain) (**Fig. 2C**), as well two novel subsets. One of these (designated B-Timm8a1+) was marked by expression of *C1qbp* (encoding a complement inhibitory protein), *Tnf*, *Il4i1*, and *Timm8a1* (encoding a mitochondrial translocase) (**Fig. 2C** and **Table S2**). The final subset (B-Uvrag+) expressed high levels of *Uvrag*, a gene involved in autophagy and tumorigenicity (**Fig. 2C**). These results reveal a diversity of ovarian B cell subsets, including a prominent presence of classic B2 cells as well as innate-like B1a and B1b populations.

**Figure 2.**
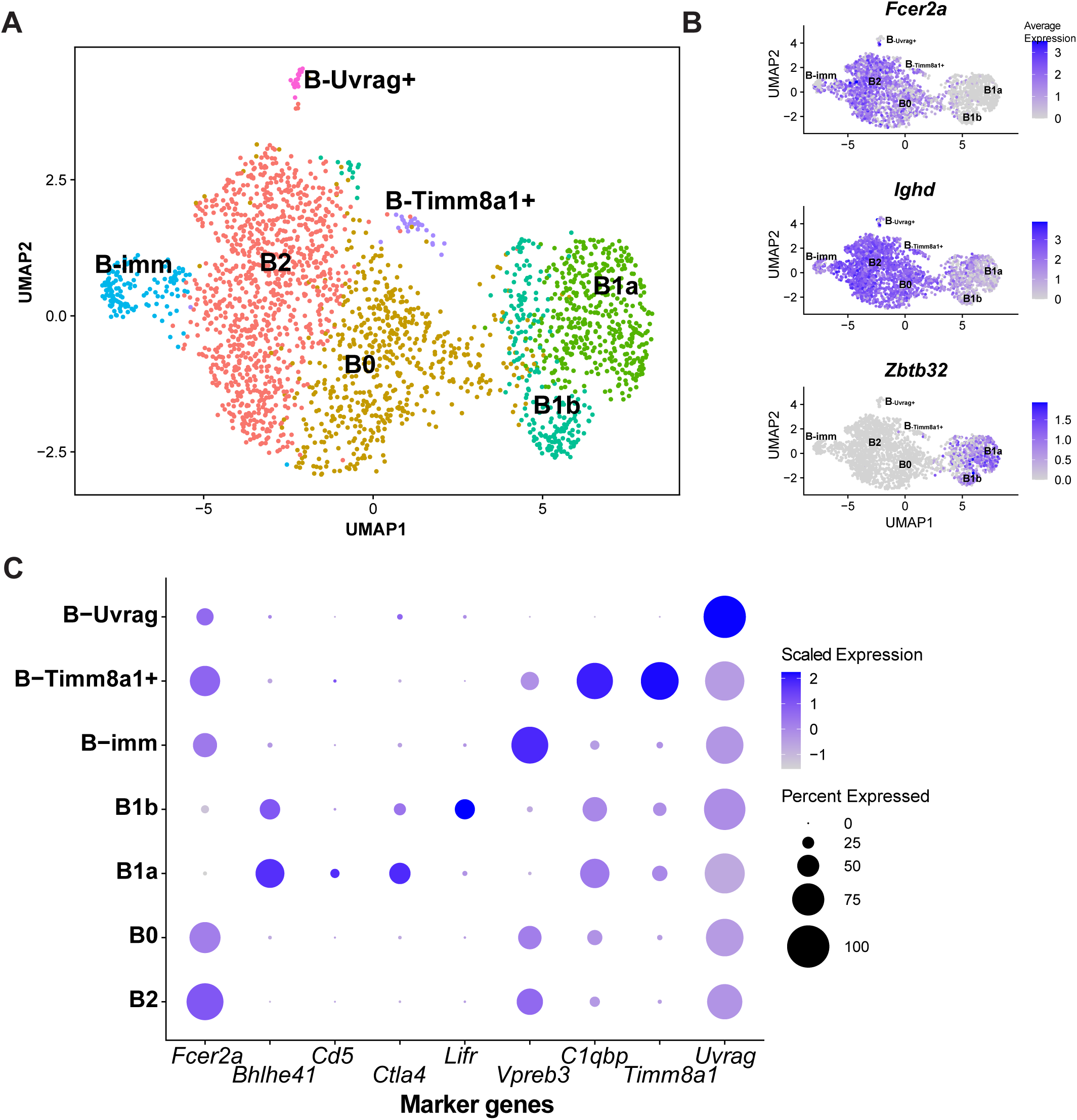
scRNAseq analysis of ovarian B cell populations. B1, B2, and B0 populations in Fig. 1 were selected for sub-clustering analysis using the Seurat package, and new cell clusters were identified. (**A**) UMAP-based two-dimensional projection of B cell subpopulations and transcriptional states. (**B**) Feature plot of selected B1 and B2 marker genes of interest. (**C**) Dot plot of scaled expression of selected marker genes for B cell subpopulations.

### Analysis of ovarian NK cells identifies mature and immature subsets

We next performed re-clustering analysis of NK cells, including the two *Ncr1+* subsets identified earlier: conventional NK cells and gd-NK cells. This analysis yielded five distinct clusters (**Fig. 3A**). Examination of known markers of mature (*Itgam*, *Klrg1*, *Gzmb*) and immature (*Cd27*) NK cells revealed a gradient of maturation, with two clusters appearing to represent the most mature populations **(Fig. 3B**). The larger of these two was designated NK-mature, while the smaller was designated NK-Ccl3-4 based on its high expression of *Ccl3* and *Ccl4* chemokines (**Fig. 3B-C**). NK-mature cells expressed abundant cytotoxic markers like *Gzma*, *Gzmb*, and *Prf1*, as well as characteristic NK receptors including *Klra8, Klra9* (Ly49 family genes), *Nkg7*, *Klrd1*, and the chemokine *Ccl5* (**Fig. 3B-C** and **Table S2**). The second-largest cluster comprised immature NK cells (NK-immat), marked by expression of *Cd27*, *Ccr2*, *Ctla2a*, and the transcription factor *Tcf7* (**Fig. 3B-C** and **Table S2**). An additional NK cells subset (NK-Srm) was distinguished by expression of *Srm* (involved in polyamine metabolism and implicated in NK cell function (28–30)), *Tsr1*, and *C1qbp*, and numerous other genes of as-yet unknown significance (**Fig. 3C** and **Table S2**). The final cluster corresponded to the previously identified gd-NK cluster and was marked by expression of *Tcrg-C4*, *Cxcr6*, and *Il7r*, among other genes (**Fig. 3C** and **Table S2**). These findings indicated the presence of both mature and immature NK cells in the ovary, with mature NK cells exhibiting strong cytotoxic potential.

**Figure 3.**
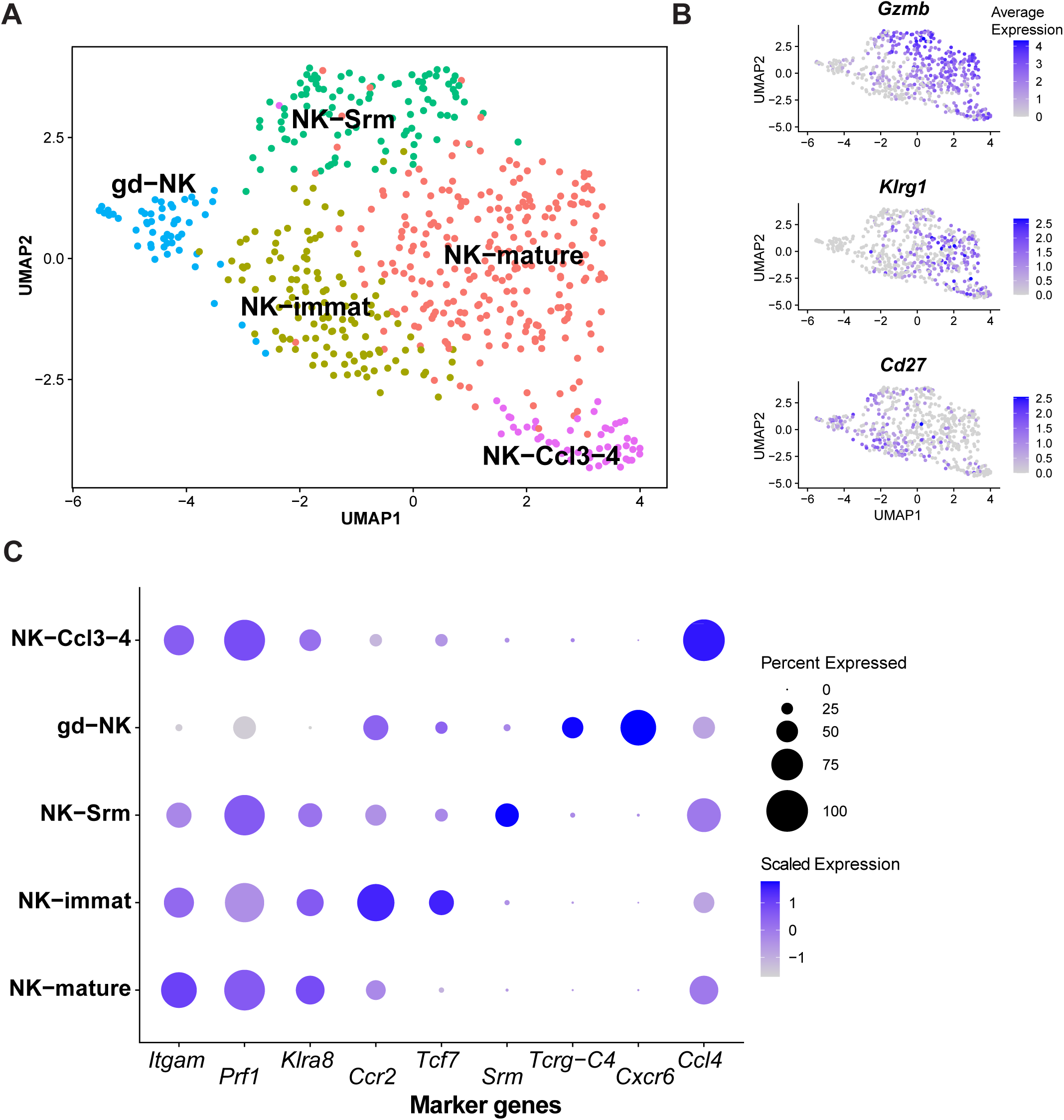
scRNAseq analysis of ovarian NK cell populations. NK cells and gd-NK populations in Fig. 1 were selected for sub-clustering analysis using the Seurat package, and new cell clusters were identified. (**A**) UMAP-based two-dimensional projection of NK cell subpopulations and transcriptional states. (**B**) Feature plot of known maturation state marker genes for NK cells. (**C**) Dot plot of expression of selected marker genes for NK cell subpopulations.

### Analysis of ovarian **α**βT cells subsets identifies Treg cells, naive CD4 and CD8 cells, and diverse mature subsets

Lastly, we performed a sub-clustering analysis on αβT cells, which included the three original clusters: CD4 T, CD8 T, and DN T cells. These predominantly expressed the αβTCR, as assessed by *Trac* expression (**Fig. 1C**). This analysis revealed seven clusters: one expressing *Cd8b1* (CD8), three expressing CD4, two mostly lacking CD4 and CD8, and one that comprised a mixed population of ∼50% of CD4 and 50% CD8 cells (**Fig. 4A**). Assessment of naive (*Ccr7* and *Sell*; encoding CD62L) vs antigen-experienced (*Cd44*) marker expression revealed a maturity gradient, with naive CD4 and CD8 T cells clustered most distinctly apart, while non-naive subsets, including DN T cells, clustered together (**Fig. 4B**). Lineage marker analysis identified discrete Th1 (*Tbx21+ Ifng+*) and Treg (*Foxp3+*) clusters (**Fig. 4C**). The Th2 marker *Gata3* was expressed at low to intermediate levels across most clusters, and Th17 cells were notably absent, as indicated by lack of *Rorc* and *Il17a* expression (**Fig. 4C**). The most abundant cluster consisted of naive CD8 T cells (CD8-naive), identified by the markers above (**Fig. 4A** and **D**). The next two largest clusters were naive CD4 T cells (*Sell+, Ccr7+ Cd44+*), distinguished by *Tcf7* and *Cd40lg* expression, with *Tcf7* possibly marking a stem-like population (CD4-stem), and *Bcl2* and *Ldlr* expression marking the CD4-Bcl2+ subset (**Fig. 4D** and **Table S2**). The Th1-like cluster, containing a mix of CD4 and CD8 T cells (designated as CD4-CD8), expressed *Tbx21, Ifng, Casp1* and *Casp4,* with the caspases being associated with proinflammatory cytokine processing(31). Two distinct DN subsets were observed. Each clustered near either CD8 or CD4 subsets, and a small fraction of those cells within them expressed *Cd8b1* and *Cd4*, respectively (**Fig. 4A**, **D**), suggesting lineage proximity. The CD8-proximal subset (DN-CD8-Ly49) expressed high levels of NK-associated marker genes, including multiple Ly49 family (*Klra1*, etc.) (**Fig. 4D** and **Table S1**), suggesting they may represent suppressive/regulatory Ly49+ CD8 T cells (32) that have downregulated CD8 expression in the ovary. The CD4-proximal DN subset (DN-CD4-CTL) expressed *Il4*, *Tbx21*, *Ifng*, *Gzmb*, and *Fasl* (**Fig. 4D**), indicating a hybrid Th1/CTL-like phenotype, along with the Th2 cytokine *Il4*. The *Foxp3+* Treg cluster expressed canonical regulatory genes including *Il2ra*, *Ikzf2*, *Ctla4*, and *Ikzf4* (**Fig. 4D, Table S2**), though *Il10* expression was not detected. This cluster did express *Tgfb1*, an immunoregulatory cytokine that was also seen across most other T cell subsets, and most prominently in DN-CD4-CTL cells (**Fig. 4C**). Taken together, these results reveal diverse αβ T cell subsets in the ovary, including abundant naive CD4+ and CD8 T+ cells, Tregs, and several unique antigen-experienced populations, such as DN/CD8-low Ly49+ T cells and DN/CD4-low cytotoxic-like T cells.

**Figure 4.**
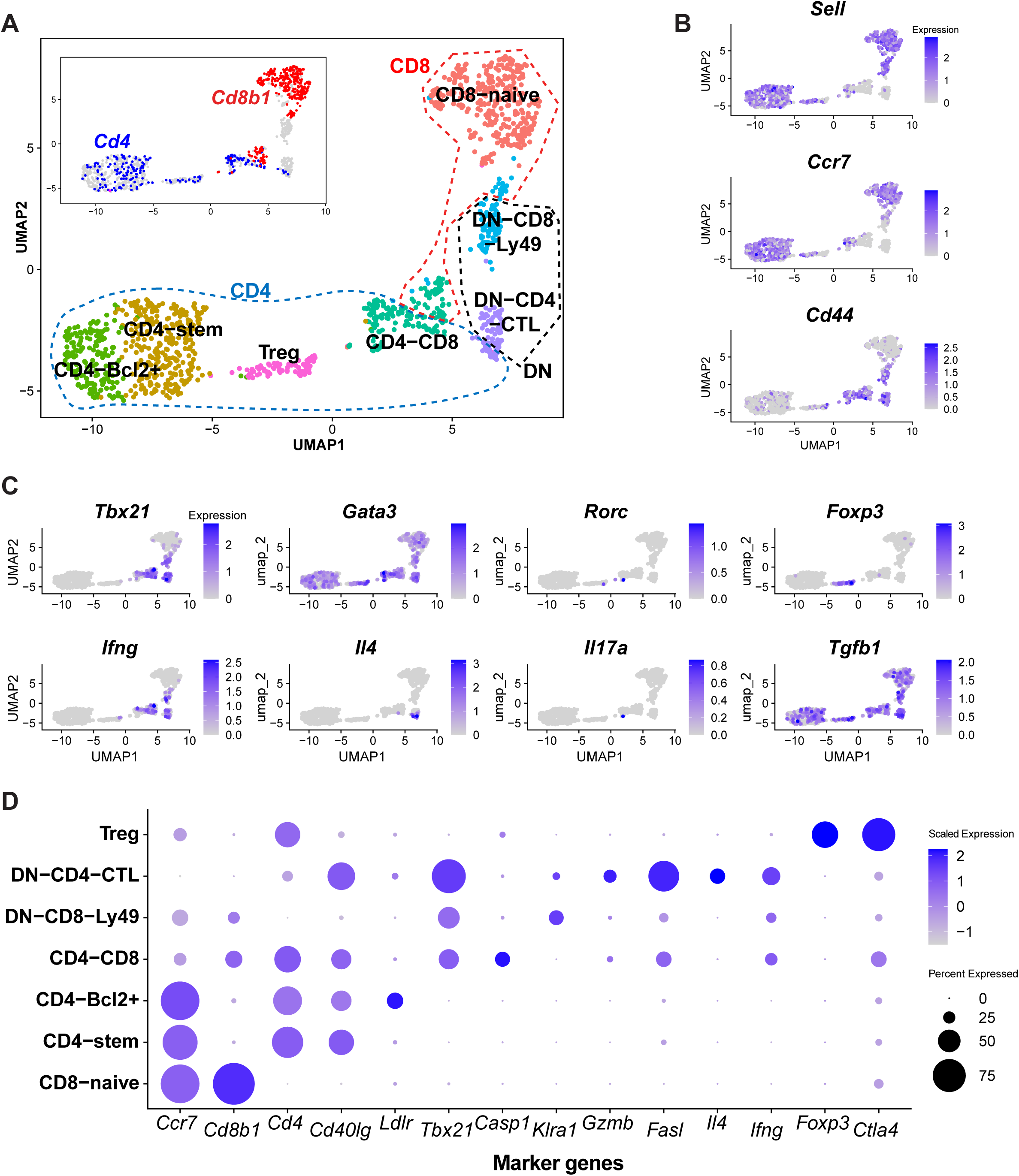
scRNAseq analysis of ovarian αβ T cell populations. αβ (i.e. Trac+) T cell populations (CD8 T, CD4 T, and DN T) populations in Fig. 1 were selected for sub-clustering analysis using the Seurat package, and new cell clusters were identified. (**A**) UMAP-based two-dimensional projection of αβ T cell subpopulations and transcriptional states. Inset shows expression of Cd4 and Cd8b1 markers. (**B**) Feature plot of known maturation state marker genes for αβ T cells. (**C**) Feature plot of known T helper cell lineage-specific transcription factors and cytokines. (**D**) Dot plot of expression of selected marker genes for different αβ T cell subpopulations.

### Ovarian immune cells are embedded in tissue and not intravascularly sequestered

We next sought to confirm that the ovarian leukocytes identified by scRNA-seq were within ovarian tissue, and not intravascular contaminants. Using a previously described technique, we i.v. injected naïve, reproductive aged C57BL/6J mice with fluorescently labeled anti-CD45 antibody (PE; CD45-i.v.). The mice were then sacrificed after three minutes, ovaries harvested, and ovarian cells were labeled with CD45 with a different fluorochrome (AlexaFluor700; CD45-ex vivo) and additional surface markers, then analyzed by flow cytometry. Peripheral blood was used as a control, and as expected all peripheral blood CD45+ cells had been intravascularly labeled *in vivo*. In the spleen a large proportion, but not all CD45+ cells had been stained *in vivo*. In the ovary, none of the CD45+ cells stained positive by intravascular CD45 labeling *in vivo*. A small contaminating CD45-i.v.+ population, likely autofluorescent cells, was identified and remained similar between the control (i.v. unlabeled) group and the intravascular labeled group. These results suggest that leukocytes identified in the ovary are ovary resident and are not intravascular contaminants **(Fig. 5)**.

**Figure 5.**
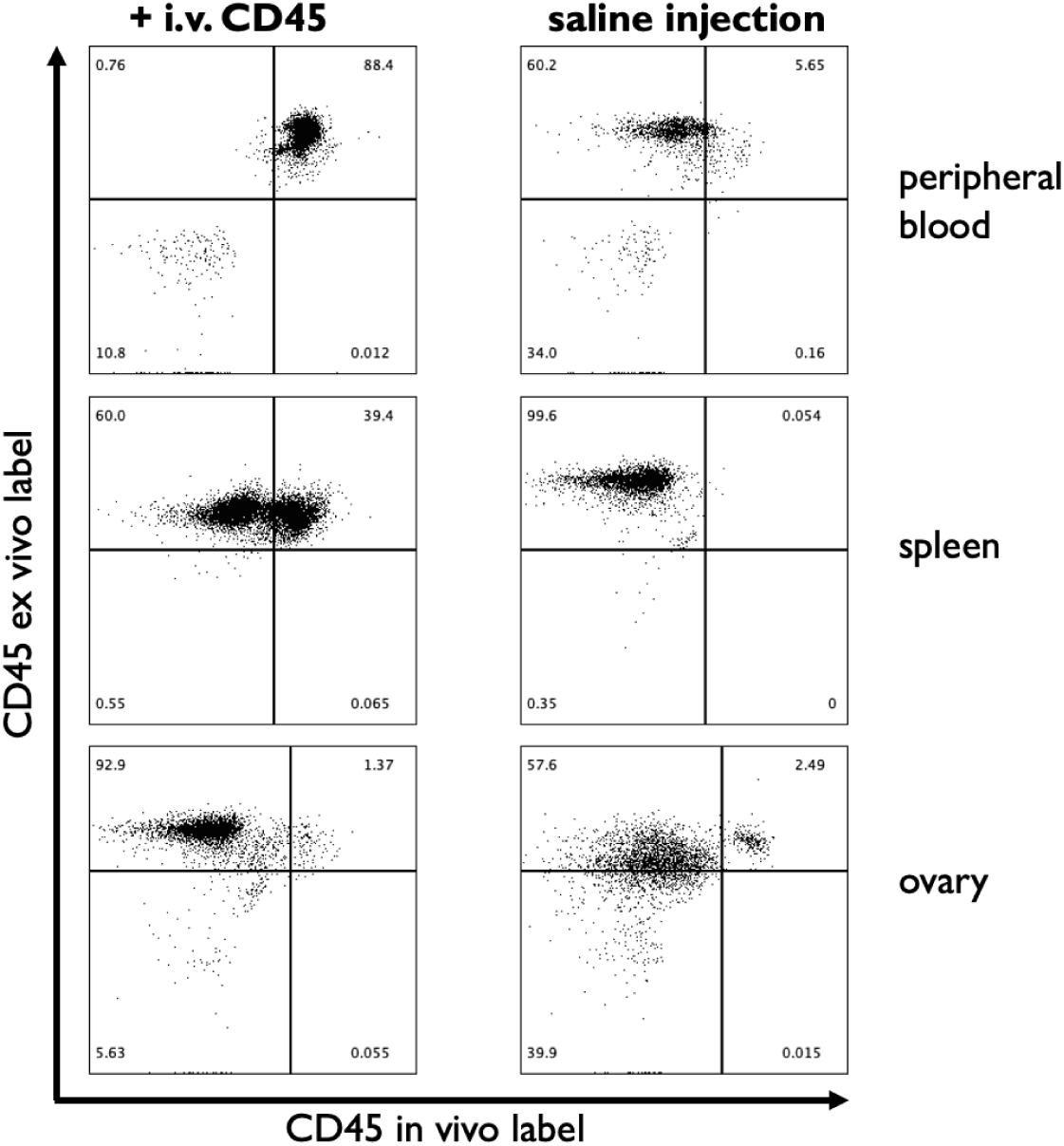
Intravascular labeling demonstrates that ovarian leukocytes are present in the tissue parenchyma. 10 week-old female C57BL/6 mice were i.v. injected with a PE-labeled anti-CD45 antibody (CD45 in vivo label) or saline control. After 3 minutes mice were euthanized and blood, spleen, and ovarian cells were collected. Cells were surface stained with antibodies directed to CD45 conjugated to AlexaFluor647 (CD45 ex vivo label) and analyzed by flow cytometry. Representative flow cytometry results for blood, spleen, and ovarian cells from i.v. labeled vs. control mice are shown. Data are representative of 2 experiments with 3-5 mice per group.

### Characterization of cell-cell communication pathways among ovarian leukocytes reveals extensive signaling by myeloid cells and NK cells, and involvement of CCL chemokines and TGFβ

A key conserved function of immune cells is cell-cell communication, predominantly mediated through secreted cytokine signals. To identify dominant signaling pathways among ovarian leukocytes, we analyzed our scRNA-seq data using the *CellChat2* package, which utilizes a well-curated ligand-receptor database to infer signaling between cell types based on expression of ligand-receptor pairs (24). Assessment of overall signaling strength by cell type revealed that 6 of the 14 identified populations, including B cells, T cells, and C1q+ macrophages, exhibited substantially higher signaling potential compared to the remaining 8 (**Fig. 6A**). The dominant signaling populations included NK and gd-NK cells as major “senders” of signals; DCs, PMN, and tissue macrophages as major “receivers”; and Mertk+ macrophages as both senders and receivers (**Fig. 6A**). Notably, many signals from NK and gd-NK cells were directed toward myeloid subsets, while signals from myeloid subsets were often autocrine or targeted to other myeloid populations (**Fig. 6B**). We next identified specific signaling pathways contributing most significantly to total signaling activity. Among the 27 pathways with significant signaling probabilities, the top four pathways (GALECTIN, CCL, MIF, and TGF-β) accounted for the majority of interactions (**Fig. 6C**). Each exhibited a distinct profile of sender and receiver cell types (**Fig. 6C**). Signaling pathway-specific analyses revealed unique patterns. CCL-family chemokine signals originated predominantly from NK and DN T cells and were received by Mertk*+* macrophages, PMN, and DCs (**Fig. 6D, left**). In contrast, TGF-β signaling was predominantly initiated by tissue macrophages and Mertk*+* macrophages and directed towards a range of leukocyte populations (**Fig. 6D, middle**). CD40 pathway signaling (top 5^th^ pathway overall) was notably more limited and originated mainly from CD4 and DN T cells, targeting myeloid cells, NK cells, and B1 cells (**Fig. 6D, right**). Together, these results identify GALECTIN, CCL, MIF, and TGF-β pathways as key mediators of intercellular communication among ovarian leukocytes. These pathways may regulate immune cell migration, spatial distribution, and trophic capacity within the ovary. Notably, TGF-β is well-established as a critical factor in ovarian homeostasis (33–35), and our findings identify ovarian myeloid cells as an abundant source of this cytokine. In contrast, the role of galectins in the ovary (outside of ovarian cancer) is not well defined and may warrant further investigation.

**Figure 6.**
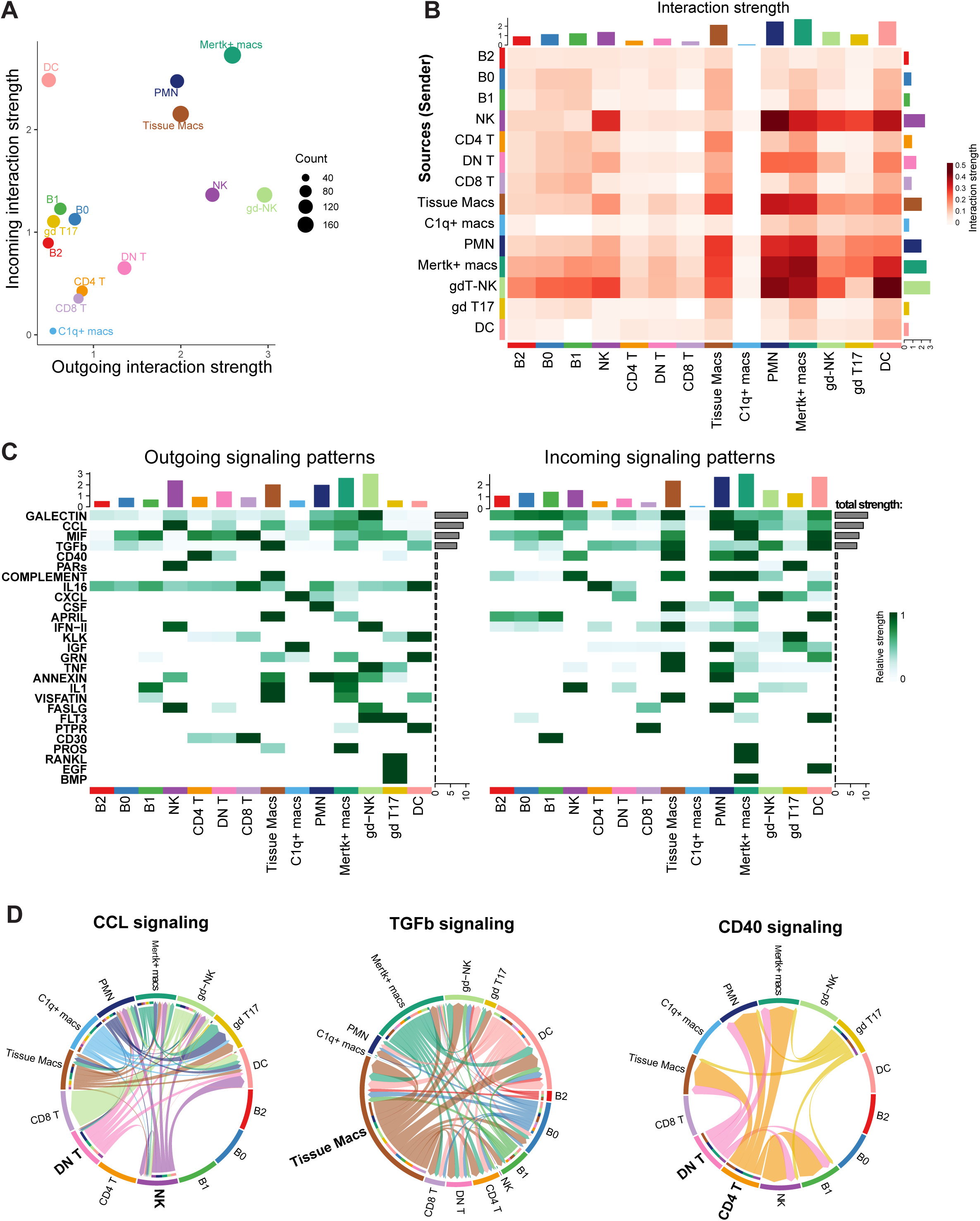
Analysis of cell-cell communication among ovarian leukocytes. Ovarian leukocyte scRNAseq data in **Fig. 1** were analyzed using the CellChat package, as described in the Materials and Methods, to infer signaling patterns among different cell subsets based on expression of secreted signaling molecules and their receptors. (**A**) Dominant signaling populations in the ovary, as inferred using incoming (Y axis) and outgoing (X axis) signal strength by population, visualized using the netAnalysis_signalingRole_scatter() function. The size of the circles indicates the total count of interactions per population. (**B**) Total signaling interaction strength between specific cell populations, visualized using the netVisual_heatmap() function. (**C**) Signaling pathway-level analysis of outgoing and incoming signals by cell population, visualized using the netAnalysis_signalingRole_heatmap() function. The grey bars to the right of the heatmap show the total signaling strength of a signaling pathway by summarizing all cell groups displayed in the heatmap per pathway. (**D**) Visualization of cell-cell signaling directionality by three selected signaling pathways (indicated above the plots), visualized using the netVisual_aggregate() function with the “chord” layout.

### Ovarian immune cells exhibit unique identities and distinct gene expression signatures

We next sought to determine how ovarian-resident leukocytes compare with those in lymphoid tissues. To this end, we used the *scANVI* package to integrate our scRNAseq data with publicly available scRNAseq data from splenocytes (see Materials and Methods). This analysis revealed that while “classic” cell types, like CD4, CD8, B2, NK cells, and tissue macrophages were abundant in both tissues, others, including B1 cells, Mertk+ and C1q+ macrophages, and gd-NK cells were unique to, or at least much more abundant in, the ovary (**Fig. 7A** and **B**). While a population similar to the one we initially designated as DN T cells was also noted in the spleen, the ovarian DN population was heterogeneous and included antigen-experienced cells, a few of which retained CD4 and CD8 expression (**Fig. 4**), which may also be the case in the splenic DN population. To identify tissue specific gene expression signatures, we performed differential gene expression analysis between ovarian leukocytes and splenocytes within each cell type. This revealed distinct gene expression signatures among ovarian leukocytes. For example: ovarian B1 cells showed higher expression of *Sox5* and *Ctla4* and lower expression of *Ifit1*; ovarian NK cells expressed more *Gpc3* and less *Irf7* and *Tnfrsf8*; and ovarian tissue macrophages expressed more *Runx2* and *Ccl7*, and less *Ccr3* and *Hebp1* (**Fig. 7C**). Pathway enrichment analysis of differentially expressed genes between ovarian leukocytes and splenocytes revealed distinct functionality. Ovarian B1 cells exhibited enhanced regulation of apoptotic pathways, while ovarian NK cells showed reduced NK cytotoxicity signatures compared to their splenic counterparts (**Fig. 7E**). Taken together, these results demonstrate that ovarian leukocytes (**A**) include unique populations largely absent from lymphoid tissues, such as C1q+ and Mertk+ macrophages and B1 cells, and (**B**) exhibit distinct tissue-specific patterns of gene expression programs that may underlie immune-regulatory functions in the ovary.

**Figure 7.**
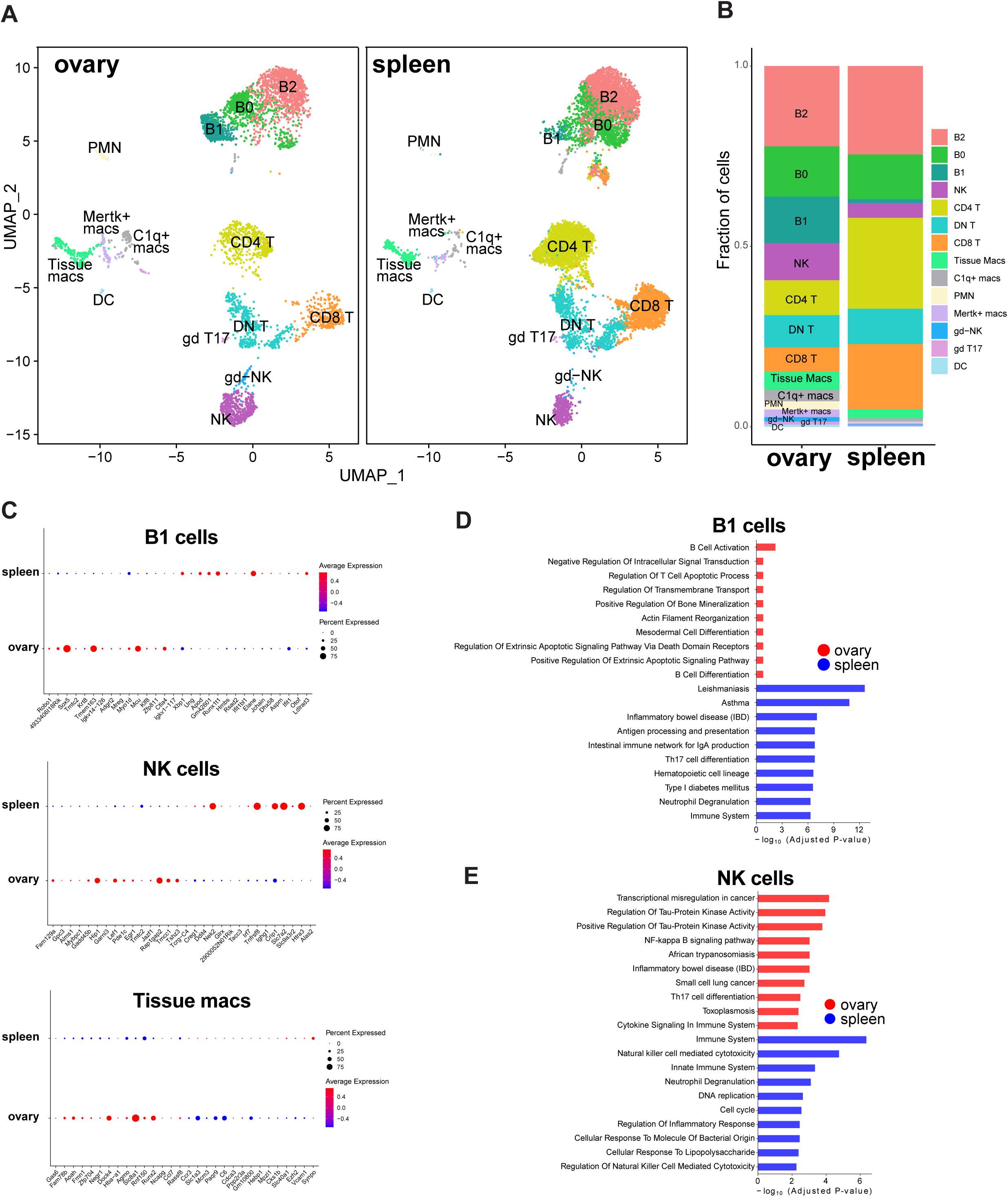
Ovarian immune cells are compositionally and transcriptionally distinct compared with splenocytes. Ovarian leukocyte scRNAseq data in Fig. 1 were integrated with publicly available data from splenocytes using the scANpy package, while maintaining the original identities of ovarian leukocyte populations. (**A**) UMAP-based projection of ovarian leukocytes and splenocytes. (**B**) Quantification of the proportion of the major cell population clusters in the ovary and spleen. (**C**) Ovarian- and spleen-specific gene expression signatures in each cell population were identified using differential gene expression analysis. Top 20 up and downregulated genes in the ovary relative to the spleen are shown for B1 cells, NK cells, and tissue macrophages. (**D**) Ovary and spleen-specific differentially expressed genes in B1 cells and NK cells were subjected to pathway enrichment analysis using gseapy v1.1.1. Top 10 enriched pathways in each tissue are shown.

### Pre-ovulatory state influences the ovarian immune microenvironment and leukocyte gene expression

Estradiol is the major hormone expressed in the ovary during the pre-ovulatory state, and ovarian-resident leukocytes are therefore exposed to substantial estradiol levels. Since the biological actions of estradiol are mediated via three receptors, estrogen receptor (ER)α, ERβ, and the membrane ER, we assessed the expression of the genes encoding these receptors (*Esr1, Esr2,* and *Gper1,* respectively) among ovarian leukocytes using our scRNA-seq data. This revealed minimal expression of *Esr2* and *Gper1*, but abundant *Esr1* expression among many, but not all, ovarian leukocytes, in particular NK and gd-NK cells, B1 cells, CD8 and CD4 T cells, as well as *Mertk+* and tissue macrophages (**Fig. 7A**). These findings suggest that ovarian leukocytes are equipped to respond to estrogen fluctuations in their microenvironment via the canonical ERα pathway.

We next used scRNA-seq to determine whether the high-estradiol state was associated with phenotypic changes in ovary-resident immune cells. Pooled data from control and GN-treated mice (**Fig. 1**) were analyzed by treatment group. UMAP-based clustering revealed abundant NK cells, B cells, CD8+ T cells, CD4+ T cells, DN T cells, and several distinct myeloid subsets in both GN-treated and control mice (**Fig. 8A**). Quantification of cluster proportions revealed multiple effects of GN treatment. In superovulated mice, the frequency of NK cells of the CD45+ compartment doubled from 7% in controls to 14% (**Figure 8B, C**). Conversely, B1 cells decreased from 18% to 8%, suggesting possible emigration and/or apoptosis in the pre-ovulation, high-estradiol state (**Fig. 8B, C**). The frequency of DN T cells and neutrophils also increased moderately but significantly in GN treated mice, while the frequency of CD8 T cells decreased (**Fig. 8B**).

**Figure 8.**
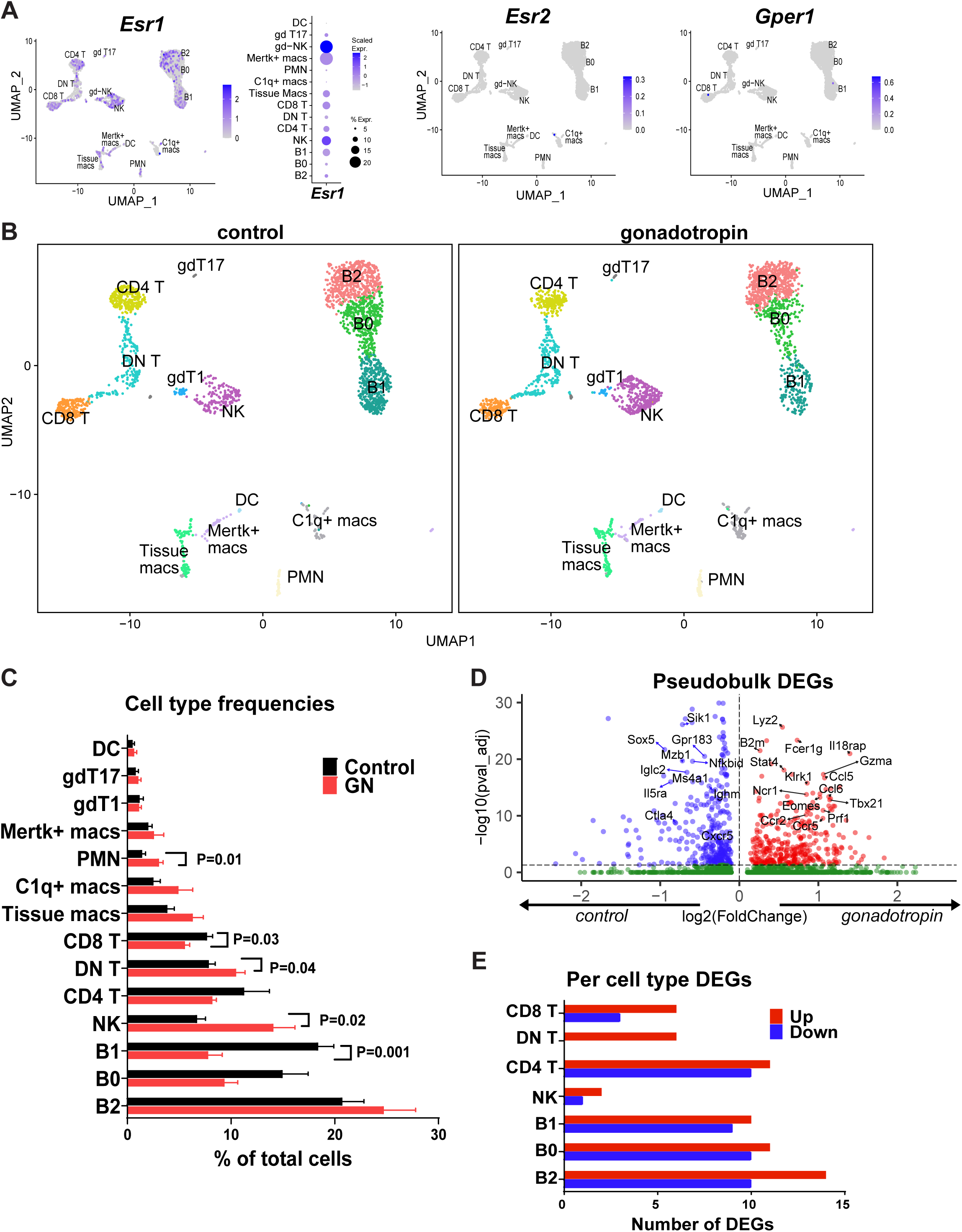
Estrogen signals driving changes in the ovarian immune landscape. Female C57BL/6 mice (n=5 per group) were injected with PBS or gonadotropin (hMG) and ovarian CD45+ cells were analyzed by scRNAseq, as described in Fig. 1. (**A**) Feature plot depicting the expression of genes encoding the major estrogen receptors, Esr1, Esr2, and Gper, among ovarian immune cells. Esr1 expression is also depicted as a dot plot. (B – E) Data presented in Fig. 1 were split by treatment (control vs. gonadotropin). (**B**) UMAP-based projection of ovarian leukocytes from control vs. gonadotropin treated mice. (**C**) Quantification of cell population frequencies in ovarian leukocytes between control and gonadotropin mice. Cell frequencies from individual mice were used for quantification, with the error bars representing the standard error of the mean. Significance was determined using a two-tailed equal variance t test. (**D**) Pseudobulk analysis of the effect of gonadotropin treatment on total ovarian leukocyte differential gene expression. Select genes of interest are labeled. (E) The effect of gonadotropin treatment on differential gene expression was assessed within each ovarian cell population. Number of up or down-regulated genes with gonadotropin treatment is shown for all populations where any significant (P_adj_<0.05) DEGs were found.

To determine whether GN treatment altered not only immune cell frequencies but also gene expression, we performed differential gene expression profiling by treatment within each cell cluster and in “pseudobulk” mode. Pseudobulk analysis revealed robust differential gene expression, largely reflecting the observed changes in cell-type proportions. For instance, NK cell genes (e.g. *Gzma, Ncr1, Klrk1*) were upregulated, while B1 cell-associated genes (*Mzb1, Ighm, Ctla4*) were downregulated in GN-treated samples (**Fig. 8D**, **Table S3**). Macrophage marker genes *Lyz2* and *Fcer1g* were also upregulated (**Fig. 8D**), consistent with a modest, albeit non-significant, increase in the frequency of tissue macrophages and C1q+ macs in the GN-treated group (**Fig. 8C**). However, when we examined differential gene expression analysis by treatment within each cell cluster, only a limited number of differentially expressed genes (DEGs) were identified (**Fig. 8E**, **Table S3**). Many of the DEGs were shared across cell types, and included cell stress-response genes such as *Jun, Fos,* and heat-shock proteins (**Table S3**), which are known artifacts of tissue dissociation in scRNAseq analysis (36, 37). Nonetheless, several unique DEGs of interest were identified, including upregulated *Stat4* in B2 cells, downregulated *Ctla4* in B1 cells, and upregulated *Foxp1* and *Satb1* in CD4 T cells (**Table S3**). The functional importance of CTLA4 in maintaining the tolerogenic functions of B1 cells is well established (38), while the latter two transcription factors have been reported to maintain quiescence or tolerance in CD4 T cells (39, 40). Collectively, these findings indicate that early estrogen-driven changes in the ovarian immune environment are characterized by shifts in cell population frequencies, with surprisingly modest transcriptional changes at the level of individual cell subsets.

Because subtle transcriptional changes can be difficult to detect across the full transcriptome, we next performed a more targeted analysis of cell-cell communication using *CellChat2*. Differential signaling analysis between GN-treated and control mice revealed that although most cell types maintained similar communication profiles, three exhibited notable changes: NK cells exhibited reduced outgoing signaling, while tissue macrophages and *Mertk+* macrophages showed increased signaling (**Fig. 9A, B**). These changes were consistent across most of the interacting cell types (**Fig. 9C**). Further pathway-specific analysis revealed that signaling through EGF, FASLG, and IFN-II pathways was stronger in controls, whereas GN-treated samples showed increased signaling through the CCL, CD40, and IGF pathways (**Fig. 9D**). A more detailed examination of specific pathways with differential signaling revealed subtle changes in the strength of specific cell-cell communication pathways (**Fig. 9E**). Taken together, these results demonstrate that early effects of superovulation include significant changes in the frequency of immune cell types and modest transcriptional changes that impact specific signaling pathways, particularly in NK cell and myeloid cell populations, which may contribute to tissue remodeling.

**Figure 9.**
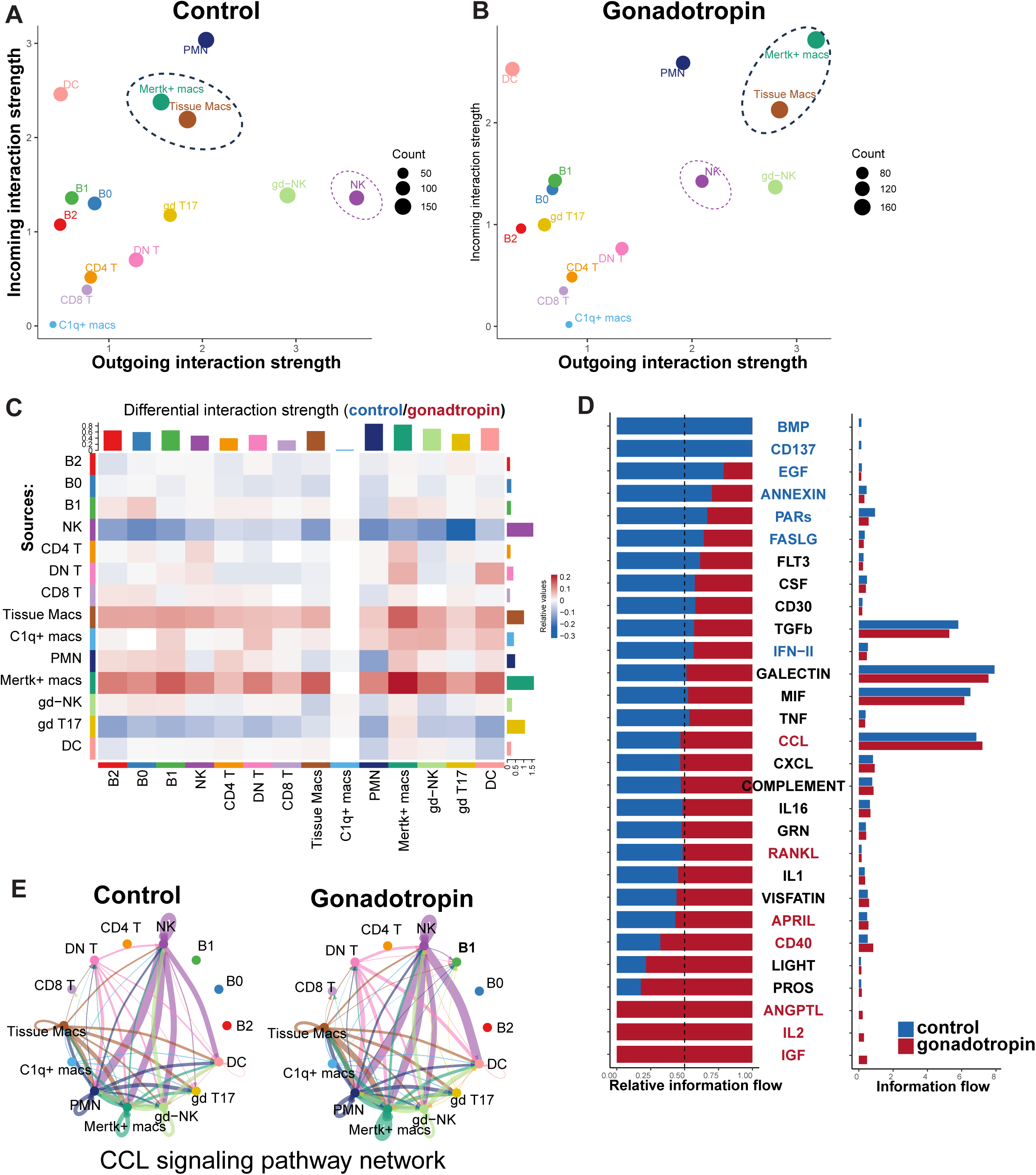
Analysis of the effect of superovulation on cell-cell communication among ovarian leukocytes. Ovarian leukocyte scRNAseq data in Fig. 8 were analyzed using the CellChat2 package for the differential effect of gonadotropin treatment, as described in the Materials and Methods. (A, B) Inference of incoming and outgoing signal strength by cell population, visualized using the netAnalysis_signalingRole_scatter() function, for control (**A**) and gonadotropin treatment (**B**) conditions. The size of the circles indicates the total number of interactions per population. (**C**) Effect of gonadotropin on signaling interaction strength between specific cell populations, visualized using the netVisual_heatmap() function, with interactions increased in control depicted as blue and those increased in gonadotropin-treated condition as red. (**D**) Specific signaling pathways showing significant differential signaling by treatment, as identified using the rankNet() function, with blue font indicating significantly higher signaling in the control condition, and red font in the gonadotropin condition (E) Visualization of cell-cell signaling directionality and networks in the CCL signaling pathway, split by treatment, visualized using the netVisual_aggregate() function with the “circle” layout.

### Validation of ovarian leukocyte transcriptional patterns during homeostasis and during ovarian stimulation at the protein level

To validate our transcriptomic findings at the protein level, we used flow cytometry to assess the composition of ovarian immune cell subsets with and without ovarian stimulation with GN. Within the ovary without stimulation, distinct populations of B cells, as assessed by CD19 staining, and αβ T cells, using TCR-β as a marker, were identified (**Fig.10A**). In addition to a small population of CD4+ and CD8+ cells, the majority of TCR-β+ cells expressed neither CD4 nor CD8, i.e. were double-negative, confirming our scRNAseq findings above. These cells did not express NK1.1, an NK cell marker, or the γδ TCR (data not shown). However, both NK and γδ T cell populations were separately identified as small populations in the CD45+/TCR-β-/CD19-gate (**Fig. 10A**). We also identified CD11b+ macrophages in the ovary (**Fig.10A**), as has been previously described (41).

**Figure 10.**
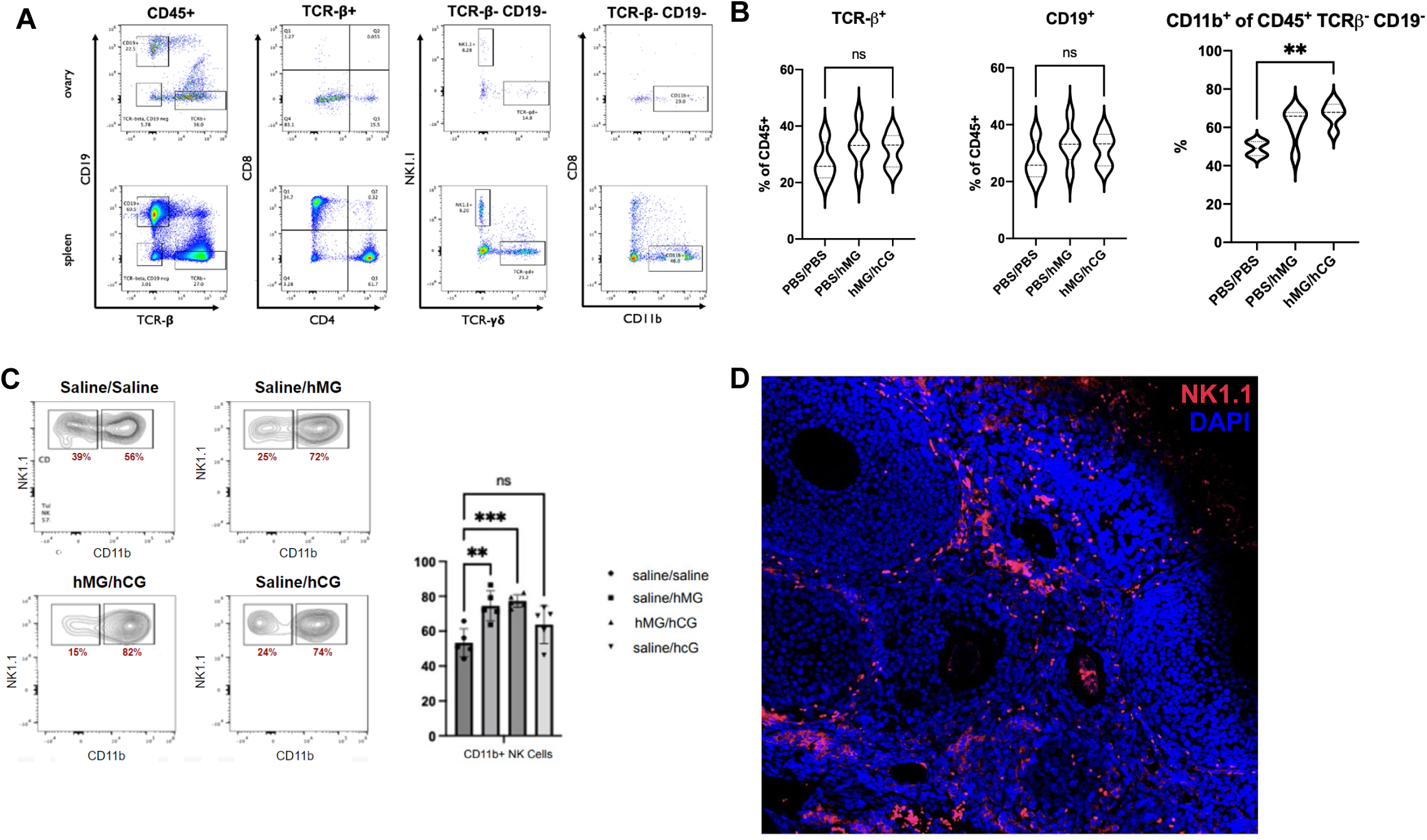
High-estrogen state influences the ovarian immune microenvironment. Ten-week-old C57BL/6 mice were injected with menopausal gonadotropins (hMG) intravenously and ovaries were harvested 48 hours later. (**A**) representative flow cytometry staining on untreated mouse ovary. (**B**) Frequencies of TCRβ+ T cells, CD19+ B cells, and CD11b+ cells, in control animals compared with high-estrogen state 48 hours after hMG injection or post-ovulation after injection with hMG and human chorionic gonadotropin (hCG). (**C**) Effect of gonadotropin treatment of frequencies CD11b+ and CD11b-populations of NK1.1+ NK cells. (D). Immunofluorescence of frozen section of untreated C57BL/6 ovary with NK1.1 in red and DAPI in blue. Representative of 4 independent experiments with 3-5 mice per group. **p < 0.01 by Student’s t-test.

Human menopausal gonadotropin (hMG) contains a combination of follicle stimulating hormone and luteinizing hormone and has been used in humans and mice to stimulate ovarian follicles, which is then followed by injection of human chorionic gonadotropin, which binds the luteinizing receptor and stimulates ovulation (42, 43). To induce both a stimulated, preovulatory peak (characterized by high estradiol, as used in the scRNA-seq experiments above) as well as a stimulated but post-ovulation phase (characterized by estradiol and progesterone), we treated with hMG alone or hMG followed by hCG, respectively. The frequencies of B cells and αβ T cells were not significantly different in hMG treated mice or post-ovulation in hMG/hCG treated mice (**Fig. 10B**). The frequency of CD11b+ cells in the CD45+/TCRβ-/CD19-gate was significantly increased in both hMG and hMG/hCG populations (**Fig. 10B**), suggesting changes in the relative abundance of macrophages or other CD11b+ cells, which could include NK cells, in response to ovarian stimulation.

Given the changes in NK frequency that we observed by scRNA-seq after GN treatment, and our findings on CD11b+ cells above, we assessed NK cell maturation state, using CD11b as a marker of mature NK cells. Mice treated with hMG or hMG/hCG, but not hCG alone, demonstrated increased frequency of CD11b+ NK1.1+ cells in the ovary (**Fig. 10C**). Flow cytometry data was confirmed with immunofluorescence, which showed abundant NK1.1+ cells palisading ovarian follicles (**Fig. 10D**). Taken together, these results confirm the presence of the ovarian major immune cell subsets identified by scRNA-seq, and demonstrate abundant mature NK cells in the ovary, which respond to estrogen flux.

## Discussion

Apoptotic debris is abundant in the ovary as part of the normal ovarian cycle, thus, the efficient clearance of apoptotic debris to prevent secondary necrosis and inflammation is presumably a well-orchestrated process, given that autoimmune POI is uncommon. Moreover, because the ovary is a dynamic tissue, it would be expected that ovarian immune cells would be similarly dynamic, but without causing chronic inflammation, which could readily lead to errors such as a break in self-tolerance. Our single-cell transcriptomic experiments revealed that the most significant population shifts after superovulation were in mature B cells and NK cells, cell types that we found were, surprisingly, the most abundant leukocytes in the ovary at baseline. The role of NK cells in normal ovary has not been described, but NK cells are implicated in ovarian cancer, and are involved in the antiviral response against mumps, a virus which is tropic to the ovary (44). Mechanistic details of these findings remain to be resolved; however, the decrease in B cells and increase in NK cells in the pre-ovulatory ovary may be related to distinct tolerance mechanisms. We propose a tripartite hypothesis whereby complex leukocyte dynamics work to prevent autoantibody production while maintaining integrity of the ovarian immune microenvironment, especially in the setting of elevated estradiol in the pre-ovulatory setting: 1) NK cell influx to provide antiviral protection with decreased threat of autoimmunity due to restricted antigen receptors, as viruses such as mumps can promote oophoritis and POI, and to promote apoptosis in stressed stromal cells that need to turnover; 2) B cell efflux to decrease likelihood of local autoimmunity as estradiol decreases tolerance threshold in B cells 3) increased signaling by tissue resident macrophages to provide trophic and tolerogenic factors like TGF-β or IGF.

Long-lived, self-replenishing B-cells, termed B1 cells produce repertoire-conserved, “natural” IgM antibodies. B1 cells arise during early development and typically reside in the peritoneal and pleural cavities (45), but can traffic to perivascular tissue (46) the spleen (45) and other tissues. Cytokines, such as IL-9 (47) IL-10 (48) IL-5 (49, 50) contribute to B1 cell development and/or function. B1 cells may play a role in recognition and removal of apoptotic (51) or senescent cells, cell debris, and other “self-antigens” generated by cellular homeostasis. With this capacity, and their production of IL-10 (52), they can help the immune system delineate non-dangerous signals and processes and prevent autoimmunity (53). However, antigen (54), cytokines e.g., IL-4 (55) or IL-5 (56), or other molecular signals, including TLR signaling (45), may also lead to harmful B1-mediated autoimmunity (48, 57–69).

The literature on B cells and NK cells throughout the estrous or menstrual cycle is lacking, and these findings contribute to our understanding of the dynamic immune microenvironment. In humans, estradiol replacement therapy has been shown to enhance peripheral blood NK cell responses, which typically wane in menopause (70). Future studies could use antibody-mediated NK or B cell depletion to assess their role in normal ovarian remodeling, as well oophoritis models. Distinct transcriptomic profiles were identified in the macrophage population, suggesting heterogeneity of ovarian tissue resident macrophages. Significant expansion or contraction of the macrophage population was not seen in GN-treated mice, suggesting stability of the tissue resident macrophages that has been described in other tissues, at least early after GN treatment. The presence of a Mertk*+* macrophage population is indicative of a population involved in efferocytosis, as would be expected given the role of this process in ovarian tissue remodeling (see above), and these cells cluster closely to *Csf1r+* tissue resident macrophages, suggesting that perhaps they represent two cell states of the same tissue resident population rather than two discrete populations. Our analysis indicates that both of these cell populations appear to play prominent role in cell-cell communication, which changes with estrogen flux, and they also serve as a source of TGFβ, a cytokine known to be important for ovarian homeostasis (33). Interestingly, we also identified an additional discrete macrophage population expressing the complement *C1q* genes, *Apoe*, and *Trem2.* This population is reminiscent of so-called disease-associated brain tissue-resident macrophages (i.e. microglia) that emerge during neuroinflammatory and neurodegenerative states (71). The presence of this population in the normal ovary suggests that in this dynamic tissue these cells may help maintain homeostasis in the face of rapid tissue turnover. Interestingly, in ovarian tumor models, blockade of TREM2 on tumor infiltrating macrophages enhances the T cell response and tumor rejection (72). Future studies can assess the role of different macrophage subsets and TREM2 in the ovary during homeostasis and inflammation.

Several recently published studies utilized scRNA-seq to assess the dynamics of the ovarian immune system in mice, predominantly focusing on the effects of aging. Isola et al (73) profiled total ovarian cells (including immune cells) from 3-month-old and 9-month-old mice. Consistent with our findings, among ovarian immune cells they identified abundant T cells, including DN T cells, B cells, NK cells, and myeloid cell populations. In their sub-clustering analyses, their results largely mirror our own, although the annotation of cell identities is somewhat different. They identified a population of *C1qa+ Apoe+* myeloid cells they designated as tissue macrophages, whereas in our own data we resolved this population as distinct from *Csfr1+* tissue macrophages. They also identified clusters designated as innate-like lymphoid cells (ILC)-1 and ILC-2, which were absent in our analysis. However, data on TCR expression by these ILC subsets was not presented, thus we suspect that they could correspond to some of the DN αβT cell subclusters, which expressed *Tbx21* and *Gata3* in our own analysis, or alternatively, NK cells, which were abundant and diverse in our studies. Interestingly, Isola et al (73) showed that aging increased the overall proportion of immune cells in the ovary, with increases in B cells, DN T cells, NKT, ILC, and γδT cells. Similar to our results, their analysis of inferred cell-cell communication also identified TGFβ as a key signaling pathway and additionally identified stromal cells and granulosa cells (absent in our experiments) as receiving TGFβ signals from T cells, macrophages, and B cells. A similar study by Ben Yaakov et al (74) used scRNAseq to analyze CD45+ ovarian cells isolated by FACS from young, middle aged, and old mice. Similar to our study and the one described above, they identified T cells, including DN T cells, NK cells, B cells, and macrophages. Like the above study, they also found cell clusters designated as ILC1s, although phenotypically they appear very similar to NK cells in our study, and we suspect that they are overlapping cell states. They found that aging decreased NK cell and ILC1 frequencies, and greatly increased the number of DN T cells, echoing some but not all of changes seen with gonadotropin treatment in our study. Lastly, Landry et al (75) used scRNA-seq to examine fibrotic changes in the aging mouse ovary. Similar to our findings, they identified three distinct macrophage subsets, which changed with aging, although their phenotypes and gene expression patterns were not described in detail. As above, several minor subsets were designated as ILCs, although they appear to be phenotypically similar to T cells and NK cells. Notably, many of the ILC populations in the above-mentioned three studies show much greater abundance in aged mice, thus it is possible that they are absent in the young adult mice used in our own study. Taken together, these scRNA-seq studies are mostly in agreement with our own findings. However, we provided a more granular analysis and annotation of various subpopulations, such as T cells (including a Treg population that was notably absent in the above studies, as well as various activated T cell populations), abundant B1a and B1b populations, and sub-populations of myeloid cells, including tissue macrophages, disease-associated-like *Apoe+* macrophages, and *Mertk+* macrophages. Together with the above studies, our findings reveal a diverse immune cell landscape in the ovary that is dynamic during both aging and ovulation, with each cell population deserving of further functional study.

